# Quantitative determination of longitudinal CNS cholesterol loss during myelin damage and repair

**DOI:** 10.64898/2026.06.04.729895

**Authors:** Disni Dedunupitiya, Eden P. Go, Travis Witte, Ashlynn Elliott, Nishama De Silva Mohotti, Jenna M. Williams, Hiroko Kobayashi, Rashmi Binjawadagi, Heather Desaire, Meredith D. Hartley

**Author notes:** **Contact information for the corresponding author:** Full Name: Meredith Hartley, Affiliation: Department of Chemistry, University of Kansas, Lawrence, Kansas, USA. Postal address: Multidisciplinary Research Building, 2030 Becker Drive, Lawrence, Kansas, 66047.

## Abstract

Cholesterol in the central nervous system (CNS) is largely unesterified (>99%) and is predominantly present in the myelin sheath (∼70% of total CNS cholesterol). Damage to the myelin sheath can result in the conversion of cholesterol to cholesterol esters, which occurs in many neurological diseases, including multiple sclerosis. In this study, we measured longitudinal CNS free cholesterol and cholesterol ester levels in a genetic mouse model during postnatal myelination, demyelination, and remyelination using gas chromatography-mass spectrometry with single ion monitoring technique (GC-MS-SIM) and liquid chromatography mass spectrometry (LC-MS). Cholesterol levels in healthy mouse brains increased up to 38 weeks. In contrast, cholesterol in the healthy spinal cord increased during postnatal timepoints, but then remained steady out to 38 weeks. Interestingly, cholesterol esters in the spinal cord were highest at P1 and drastically reduced by P42, while the brain had similar levels during all postnatal time points. During demyelination, both brain and spinal cord cholesterol levels were significantly reduced as compared to healthy mice and failed to return to normal cholesterol levels even during remyelination. Absolute quantification of cholesterol esters during peak demyelination revealed that cholesterol esters comprise 19% of the total cholesterol pool in the brain and 65% in the spinal cord. The lack of recovery in CNS cholesterol levels after demyelination suggests that healthy *de novo* cholesterol synthesis pathways are disrupted in this model. Absolute quantification of CNS cholesterol is critical for revealing mechanisms of cholesterol regulation during disease and identifying targets for restoring cholesterol to promote myelin repair.

## 2. INTRODUCTION

Cholesterol plays critical structural, biophysical, and metabolic roles in the central nervous system (CNS). The CNS tissue contains approximately 20% of the whole-body free cholesterol pool, although it comprises only 2% of the body mass (1, 2). *De novo* synthesis of cholesterol is the primary source of cholesterol in the CNS, and the blood-brain barrier blocks entry of peripheral cholesterol into the brain under normal physiological conditions. Brain cholesterol is a structural component of myelin (∼70 % of total brain cholesterol) and of glial and neuronal cell membranes (∼30%) (3). The myelin sheath is a lipid-rich multilamellar structure that wraps segments of neurons, improving the efficiency of the rapid conduction of nerve impulses. Myelin contains a higher percentage of lipids (∼80% lipids, 20% proteins) relative to typical cellular membranes (∼40% lipids, 60% proteins) (4, 5). The higher proportion of lipids in myelin causes a more tightly packed arrangement of the membrane than in regular biological membranes (6).

Myelin is synthesized by oligodendrocytes in the CNS, and cholesterol biosynthesis in oligodendrocytes is a rate-limiting factor in myelin membrane growth during brain development and maturation. Mice that lack cholesterol synthesis in myelinating oligodendrocytes showed severely perturbed myelination and suffered from ataxia and tremor (7). Most cholesterol in the brain is unesterified, with <1% of total cholesterol existing in the esterified form as cholesterol esters (8). Cholesterol comprises ∼40% of myelin lipids and is a key structural component of the myelin membrane that contributes to myelin membrane stability (9). Repaired myelin showed reduced biophysical stability relative to healthy myelin, which was due in part to reduced cholesterol levels in repaired myelin (10). Cholesterol also plays a critical role as a metabolic precursor for neuroactive steroids, which regulate myelin formation, regeneration and neuroprotection (11).

Disruption of CNS cholesterol homeostasis has been observed in many neurological diseases, suggesting an interconnection between neuronal health and cholesterol (12). Clinical studies often measure cholesterol in serum or plasma, but peripheral levels of cholesterol do not reflect CNS levels of cholesterol due to the blood brain barrier, and studies on cholesterol levels in CNS are limited due to tissue accessibility. In Alzheimer’s disease, metabolomic and transcriptomic data showed abnormal brain cholesterol biosynthesis and catabolism (13), and reduced CNS cholesterol levels inhibited the formation of Aβ plaques in primary rat hippocampal neurons (14). Evidence of disruption of cholesterol biosynthesis has also been observed in mouse models and human postmortem tissues in Huntington’s disease (15), a neural cholesterol accumulation in early Niemann-Pick type C1 (NPC) mouse brain (16), and Smith-Lemli Opitz syndrome (17). Aging is also considered a form of cognitive decline, characterized by altered short-term memory and learning, and a decline in cholesterol synthesis rate was observed in both the human hippocampus (18, 19) and rodent hippocampus (20).

Due to the high levels of cholesterol in myelin, diseases featuring loss of myelin have been linked to cholesterol metabolism and regulation. The most prevalent neurodegenerative disease with primary myelin damage is multiple sclerosis (MS), in which inflammatory demyelination leads to degeneration of neurons in the CNS (21). Blocking cholesterol synthesis in either oligodendrocytes (7) or neurons (22) greatly impaired remyelination in mouse models of demyelination underscoring the requirement of cholesterol for myelin repair. The conversion of cholesterol to cholesterol ester is associated with myelin damage and has been observed in brain white matter, spinal cord, and cerebrospinal fluid of people who were diagnosed with multiple sclerosis (23–25). Elevated cholesterol esters during demyelination have also been observed in preclinical demyelination models including the cuprizone model (26), experimental autoimmune encephalomyelitis (EAE) model (27), and a genetic mouse model of demyelination (28). Thus, robust, quantitative methods are needed to measure both cholesterol and cholesterol esters in the CNS during both development and disease.

In our study, we profiled the levels of free cholesterol and cholesterol esters during development and during demyelination and remyelination in a genetic mouse model using robust, internal standard-based quantification methods. We observed steady increases in free cholesterol levels in both the brain and the spinal cord during development. Cholesterol ester levels were highest at P1 in the spinal cord and decreased rapidly up to P42, whereas the developmental brain had similar levels of cholesterol esters during the same period. Following healthy mice out to 38 weeks, cholesterol levels continued to increase in the brain but plateaued in the spinal cord. When demyelination was induced in adult mice, cholesterol levels in the brain and spinal cord dropped and did not recover to healthy levels even after the damaged myelin was repaired. Significant accumulation of cholesterol esters was observed during demyelination indicating a shift in the overall pool towards the esterified state. Our results suggest that demyelination inhibits normal cholesterol biosynthesis in this model resulting in reduced CNS levels of total free cholesterol. Quantitative determination of CNS cholesterol is critical for revealing mechanistic insights into how cholesterol metabolism is related to pathophysiology of neurological diseases that may enable the discovery of new therapeutic targets for disease.

## 3. MATERIALS AND METHODS

### Animal Studies

A genetic mouse model of demyelination with a C57BL6/J background was utilized in this study where *Myrf(fl/fl)* mice were crossed with *Plp1*-*CreERT* mice to generate the *Plp1-CreERT; Myrf (fl/fl)* strain (or *Plp1*-iCKO*-Myrf)* at the University of Kansas (29–31). At 8 weeks of age, mice were administered with intraperitoneal injections of tamoxifen for five days (100 µL of 20 mg/mL daily in corn oil) to induce *Myrf* ablation and demyelination. Mice with *CreERT* (or *Cre*-positive) lose *Myrf* in oligodendrocytes and undergo demyelination, and those without *CreERT* (or *Cre*-negative) serve as healthy controls. Mice were randomly assigned to five time points and euthanized (carbon dioxide inhalation and cervical dislocation) at 6, 12, 18, 24, and 30 weeks post-tamoxifen injection. The brain and spinal cord tissues were collected and stored at −80 ℃. Additionally, a randomly chosen cohort of *Cre*-positive and *Cre-*negative mice from weeks 12 and 18 post-tamoxifen weeks received Sob-AM2 chow treatment starting 2 weeks post-tamoxifen. Sob-AM2 was compounded into chow (Envigo Teklad 2016 diet) at a dose of 420 µg/ kg chow (a nominal daily dose of 84 µg/ kg body weight). All procedures were approved by the University of Kansas Institutional Animal Care and Use Committee.

### Modified Bligh & Dyer lipid extraction protocol for the brain homogenates of healthy and demyelinating mice

Brain tissues were homogenized (Bead Mill Homogenizer, OMNI International, 19-2241E) at 50 mg/mL or 300 mg/mL (10 ceramic beads (OMNI International, 19-645-3) at a speed of 3.10 m/s for 3 cycles of 20 s including a 1 s dwell time. Spinal cord tissue was homogenized at 65 mg/mL (3 ceramic beads, 4.0 m/s, 2 cycles of 20 s, 1 s dwell time) in ice-cold micro pure water. The brain and spinal cord tissue homogenates were aliquoted and stored in −80 ℃.

Due to the large number, samples were randomized into five batches (n = 15-20 in each batch) to perform modified Bligh & Dyer lipid extraction (32). Brain homogenates (6 mg tissue total amount) were diluted to 6 mg/mL in 1 mL of ice-cold micro pure water. Each sample was vortexed well, and 10 µL from each sample was saved for protein quantification using BCA assay. The remaining 990 µL homogenate was added into glass tubes (VWR International, 72690-022) containing CHCl_3_ (with 0.01% butylated hydroxytoluene, BHT): MeOH: water in 3:2:1 ratio. Then 5 µL of 2.54 mM d_7_-cholesterol in CHCl_3_ (Cayman Chemicals, 25265) was spiked into each tube. The glass tubes were shaken and vortexed to allow mixing. Then, 2.5 mL of CHCl_3_ was added to each glass tube, followed by the addition of 1.25 mL of water. Again, the glass tubes were shaken and vortexed ten times and five times, respectively. The samples were centrifuged (ThermoScientific Sorvall ST40R, 42508141) at 400 x g for 10 minutes at 25 ℃. Then the lower CHCl_3_ layer was removed into a small glass tube (VWR International, 72690-024), and the extractions were repeated with another 1.25 mL of CHCl_3_ for two more times. All three extracted CHCl_3_ layers were combined and washed with 300 µL of 1 M KCl and 300 μL of water, respectively (centrifuged at 400 x g for 5 minutes at 25 °C). The CHCl_3_ layer was vacuum dried (ThermoScientific Savant, SPD130DLX). Then 0.5 mL of CHCl_3_ was used to resuspend the lipids and transferred to PTFE-capped glass vials for storage. Finally, the samples were vacuum dried again, weighed, and stored in the −80 ℃ freezer. Each extracted lipid droplet was resuspended in 100 µL and 10 µL (10%) was used for the derivatization reaction.

### Modified Bligh & Dyer lipid extraction protocol for spinal cord homogenates of the healthy and demyelinating mice

Spinal cord samples were also randomized (four batches) prior to the lipid extraction. The modified Bligh and Dyer protocol as described above was followed for the spinal cord homogenates with the following exceptions: spinal cord homogenates (3.25 mg tissue total amount) were diluted to 3.25 mg/mL in 1 mL of ice-cold micro pure water, and 100 µL was saved for the BCA assay. The remaining 900 µL was used for the extraction, and 10 µL of d_7_-cholesterol (1 mM standard) was spiked in each tube. Each extracted lipid droplet was resuspended in 100 µL and 20 µL (20%) was used for the derivatization reaction.

### Lipid extraction protocol for postnatal mouse brain and spinal cord homogenates

The same lipid extraction protocol was followed for the postnatal mouse brain (5 mg of brain tissue in 1 mL ice-cold micro pure water) and spinal cord (2.5 mg tissue in 1 mL ice-cold micro pure water). The spiked amounts of d_7_-cholesterol (1 mM standard) were 15 µL and 20 µL for the brain and spinal cord, respectively. Each extracted lipid droplet was resuspended in 100 µL and 20 µL (20%) was used for the derivatization reaction.

### Sample derivatization for the GC-MS

Extracted healthy and demyelinating lipid droplets (both brain and spinal cord) were resuspended in 100 µL CHCl_3_ (0.01% BHT) and sonicated (Branson M2800) for 15 minutes. Then, respective volumes of each sample (as described above) were transferred into reaction vials. Gas-tight syringes (Hamilton) were used for transferring all organic solvents throughout the protocol. The reaction vial wall was washed with 300 µL of CHCl_3,_ and then the CHCl_3_ was vacuum evaporated. Next, 60 µL of N, O-bis(trimethylsilyl)trifluoroacetamide with trimethylchlorosilane (BSTFA + TMCS, 99:1, v/v) derivatizing reagent (LiChropur, 15238) was added and incubated at 50 °C for 1 hour. Samples were cooled to room temperature, and 140 µL of n-hexane was added. Then the reaction mixtures were vortexed and transferred into glass vials (ThermoScientific, C4013-2) with glass inserts (Supelco, 24701) and 1 µL of each mixture was injected into the GC-MS. (Note: The trimethyl silyl derivatized cholesterol can degrade after 48 hours at room temperature. Therefore, all the experiments for each batch were performed over two days following the same timeline)

### Preparation of calibration curves

Standard solutions of d_0_-cholesterol (0.50 mM) and d_7_-cholesterol (2.54 mM) were prepared using the solvent CHCl_3_ (0.01% BHT). BSTFA + TMCS, 99:1 v/v, reagent was used to derivatize standards following the protocol described above. Derivatized standards were dissolved in n-hexane before GC-MS injections. Separate calibration curves were obtained for each batch of samples. For the brain samples calibration curve, cholesterol standards were in the 4.0 – 80.0 µM range with a d_7_ cholesterol spike of 12.70 µM while spinal cord calibration curve samples were in the 10.0 µM - 0.20 mM range with a d_7_ spike of 40.0 µM.

### Quality controls

Quality controls (QCs) were prepared for the brain and spinal cord by mixing equal volumes of representative samples selected from each time point, including both *Cre*-positive and *Cre*-negative mouse groups. The mixtures were vortexed well and diluted to a final concentration of 50 mg/mL for brain samples and 65 mg/mL for spinal cord samples. The quality control mixtures were aliquoted, and three aliquots per batch were extracted, derivatized and analyzed by GC-MS alongside the other samples in the batch. Two to three quality controls were incorporated for each batch.

### Gas chromatography-mass spectrometry (GC-MS) conditions and data analysis

Gas chromatography-mass spectrometry was performed using a capillary SH-I-5MS Column (0.25 mm, 0.25 µm film, 30.0 m) on a Shimadzu QP2010 SE (Columbia, MD). Helium was used as a carrier gas at a flowrate of 1 mL/min. The initial column oven temperature was held at 150 ℃ and the injection temperature was set to 290 ℃. The initial temperature of 150 ℃ was ramped to 305 ℃ at 30 ℃/ min and then ramped at 1 ℃/ min to 315 ℃ with a hold of 1 min at 315 ℃. The total run time was 17.17 minutes. Electron impact ionization was performed at 70 eV with ion source temperature of 230 ℃ and an interface temperature of 280 ℃. The sample was injected at a volume of 1 µL in splitless mode. Blank samples were included between runs to ensure no sample carryover. GC-MS detection was set to single ion monitoring mode. Reproducible fragments (*m/z* 329 and *m/z* 336) were selected for the quantification of d_0_ cholesterol and d_7_ cholesterol, respectively (Figure 1A) (33). Peak areas of extracted ion chromatograms were automatically integrated using post-run Lab Solutions Software (Shimadzu).

**Figure 1:**
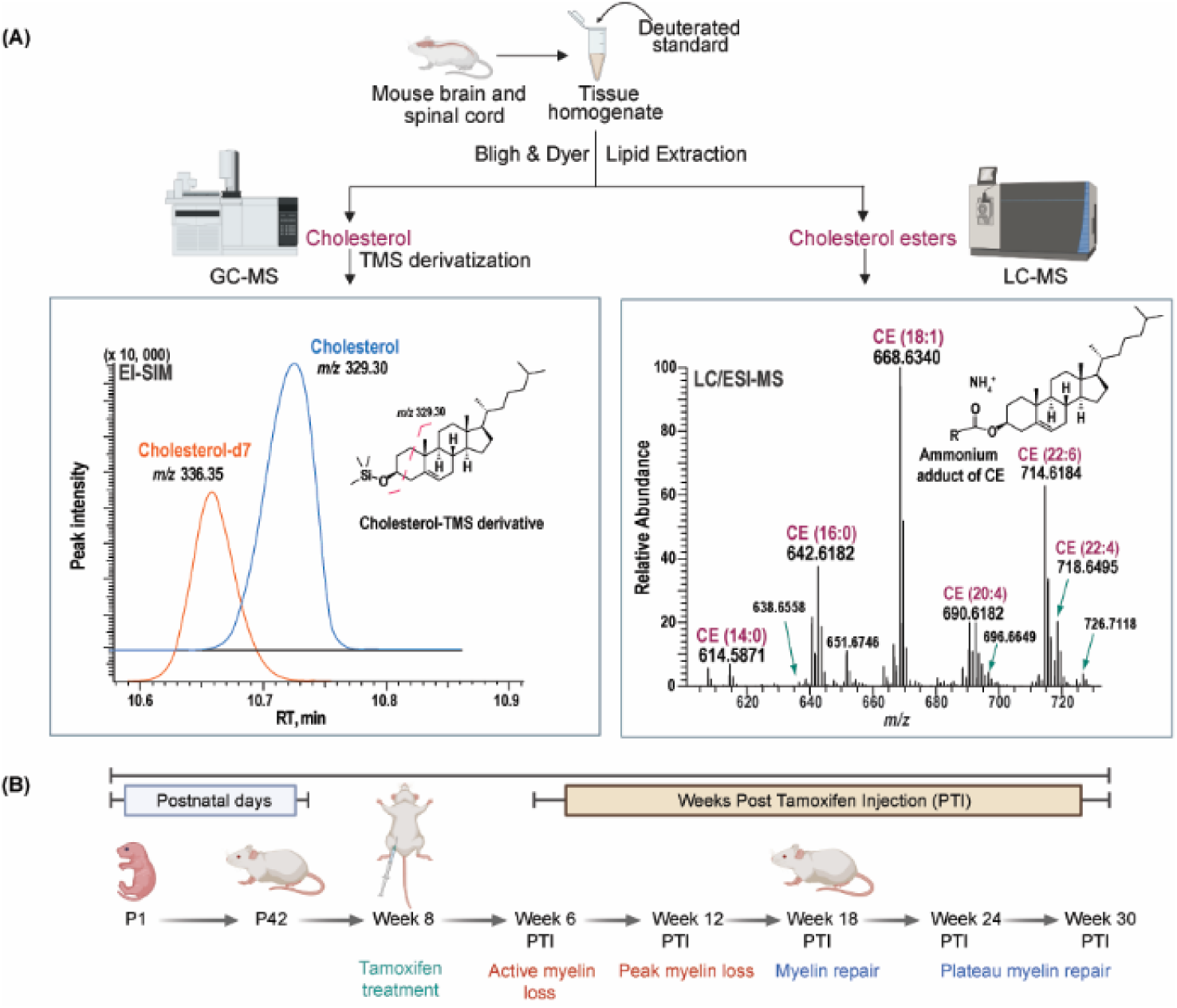
Overview of the experimental workflow for cholesterol analysis. **(A)** Experimental workflow for the quantification of cholesterol and cholesterol esters (CE) using GC-MS and LC-MS. **(B)** Timeline of the mouse model.

### Lipid extraction and liquid chromatography mass spectrometry protocol for cholesterol ester levels in mouse brain and spinal cord

To measure cholesterol esters in developmental tissues, 15 mg of developmental brain tissue and 5 mg of developmental spinal cord tissue were diluted in 1 mL of ice-cold micro pure water and the modified Bligh and Dyer lipid extraction was followed. The developmental brain and spinal cord tissues were spiked with 15 μL and 20 μL of 200 μM d_9_-cholesterol palmitate (Cayman, 28123), respectively. Brain lipid droplet was resuspended in 300 μL of 2MeOH: CHCl_3_. The resuspended lipid droplet was diluted 2.5 times (100 μL was taken out, evaporated and resuspended in 250 μL) for LC-MS injection. Spinal cord lipid droplet was dissolved 100 μL of 2MeOH: CHCl_3_ and diluted 1.5 times (100 μL was taken out, evaporated and resuspended in 150 μL) for the injection. A calibration curve was prepared using a range of 2.00 - 26.25 μM of d_0_-cholesterol palmitate (Cayman, 26473), each containing a 7.5 μM spike of d_9_-cholesterol palmitate. To measure cholesterol esters in adult CNS tissues, 15 mg of adult healthy and demyelinated brain tissue, 6.5 mg of healthy spinal cord and 3.9 mg of demyelinated spinal cord were used in diluted 1 mL ice-cold micro pure water. Modified Bligh and Dyer lipid extraction was performed with 200 μM d_9_-cholesterol palmitate spike for healthy and demyelinated brain tissue (40 μL), healthy spinal cord tissue (10 μL) and demyelinated spinal cord tissue (40 μL) respectively. Each lipid droplet was resuspended in 300 μL of 2:1 MeOH: CHCl_3_ and diluted 10 times (20 μL was taken out, evaporated and resuspended in 200 μL) for LC-MS injection. Calibration curves were prepared for adult brain and spinal cord batches in the range of 2.00 −26.25 μM of d_0_-cholesterol palmitate (Cayman, 26473), each containing a 7.5 μM spike of d_9_-cholesterol palmitate. High resolution LC-MS experiments were performed using an Orbitrap Fusion (Milford, MA). Mobile phases consisted of solvent A: ACN/H_2_O (40:60, v/v) and solvent B: H_2_O/ACN/IPA (1:9:90, v/v) both containing 10 mM ammonium formate. Five microliters of each sample were injected onto a T3 HSS C18 (3.5 μm, 1 mm, 150 mm) at a flow rate of 50 μL/min. The lipids were separated using a 15-min gradient starting at 30% B, then a 1.5 min increase to 65% B, then ramped to 99% B in 1.5 min, held for 2.0 min, and then returned to initial conditions (30% B) for 6 min. The column was re-equilibrated at initial conditions for 4 min prior to each new injection. A short wash and a blank run were performed between every sample to ensure no sample carryover. Mass spectrometric analysis was performed in the positive ion mode using the following parameters: a spray voltage of 3.0 kV, sheath gas: 10 units, auxiliary gas: 6 units, ion transfer tube maintained at 250°C, and the vaporizer set to 60°C. Full MS scans in the Orbitrap were performed in 1.5 s cycles in the mass range of *m/z* 300-1200 at a resolution of 120,000 at 200 *m/z* with an automatic gain control target value set to 4 × 10^5^ and with a maximum injection time of 50 ms. Peak intensities of ammonium adducts of twenty-one cholesterol ester types (Supp. Tables S7-S12) were manually integrated using Xcalibur software.

### Statistics and Data Analysis

Internal standard-based calibration curves were used to calculate the cholesterol level in both brain and spinal cord tissues. Cholesterol levels were normalized to tissue weight for both brain and spinal cord. Total cholesterol levels in the P1-P42 developmental time points were compared using ordinary one-way ANOVA, and Dunnett correction for multiple comparison data was performed. Data is plotted as mean ± SEM. P value is adjusted for multiple comparisons with a 95% confidence interval where 0.1234 (ns), 0.0332 (*), 0.0021 (**), 0.0002(***), < 0.0001(***). Total cholesterol esters in the developmental mouse P1-P42 timepoints using ordinary one way-ANOVA using Tukey correction for multiple comparison was used and data are plotted as mean ± SEM, GP: 0.123 (ns), 0.0332 (*), 0.0021 (**), 0.0002 (**), 0.0001 (***). Total free cholesterol levels in the *Plp* mouse brain and spinal cord were compared using Two way-ANOVA and Holm-Šídák correction for multiple comparisons. Free cholesterol data are plotted as mean ± SEM, GP: 0.123 (ns), 0.0332 (*), 0.0021 (**), 0.0002 (**), 0.0001 (***). Effect of the Sob-AM2 drug on cholesterol levels in the spinal cord. Two-way ANOVA (mixed models, Tukey correction for multiple comparisons) and the data are plotted as mean ± SEM, GP: 0.123 (ns), 0.0332 (*), 0.0021 (**), 0.0002 (**), 0.0001 (***). Total amounts of cholesterol ester (CE) and individual CE types are included in heat maps as average CE nmol amount per mg tissue. Statistical analysis was performed using GraphPad Prism 11 and figures were prepared using BioRender, GraphPad Prism 11, and Adobe Illustrator.

## 4. RESULTS

### MS methods for detecting cholesterol and cholesterol ester levels in CNS tissue homogenate samples

To accurately measure cholesterol in CNS tissue samples, we developed a quantitative method with internal standards to measure cholesterol using GC-MS with electron ionization and single ion monitoring (33, 34). In brief, the method involves collection of brain and spinal cords, tissue homogenization, lipid extraction, lipid droplet derivatization, and GC-MS sample analysis (Figure 1A). Each sample was spiked with deuterated cholesterol (d_7_-cholesterol) at the beginning of the lipid extraction, which enables more accurate quantification of the cholesterol levels in the tissues.

To derivatize the cholesterol prior to GC-MS analysis, N,O-Bis(trimethylsilyl)trifluoroacetamide with trimethylchlorosilane (BSTFA + TMCS, 99:1, v/v) was used to modify the hydroxyl of cholesterol with trimethylsilyl (TMS) to increase the volatility of the molecule. TMS-derivatization was chosen over other derivatization methods because the reaction was straightforward and generated more volatile and less polar derivatives with structurally informative fragments (35). The TMS-derivative of cholesterol ionized by EI (electron ionization) and yielded highly abundant, reproducible fragments (*m/z* 329*, m/z* 336) of cholesterol and d_7_ cholesterol, respectively, which were used for quantification (Figure 1A).

Cholesterol ester levels were quantified using ultra-high-pressure liquid chromatography electrospray ionization mass spectrometry (UPLC-ESI-MS) (36). Unlike cholesterol, cholesterol esters form stable ammonium adduct ions in the ESI source and can be detected by UPLC-ESI-MS (Figure 1A).

### Trends of free cholesterol and cholesterol ester levels in the developmental brain and spinal cord

To determine the physiological levels of free cholesterol during CNS development, we collected brain and spinal cord tissues at postnatal days P1, P4, P7, P10, P14, P21, and P42 (Figure 1B). In the brain, the cholesterol concentration was 7 nmol/mg tissue at P1 and increased to 23 nmol/mg at P42, a ∼3.4-fold increase (Figure 2A, Supp. Table S1). The spinal cord tissue showed a more dramatic increase in cholesterol starting at 12 nmol/mg at P1 and increasing to 99 nmol/mg by P42, a ∼8.1-fold increase (Figure 2B, Supp. Table S2).

**Figure 2:**
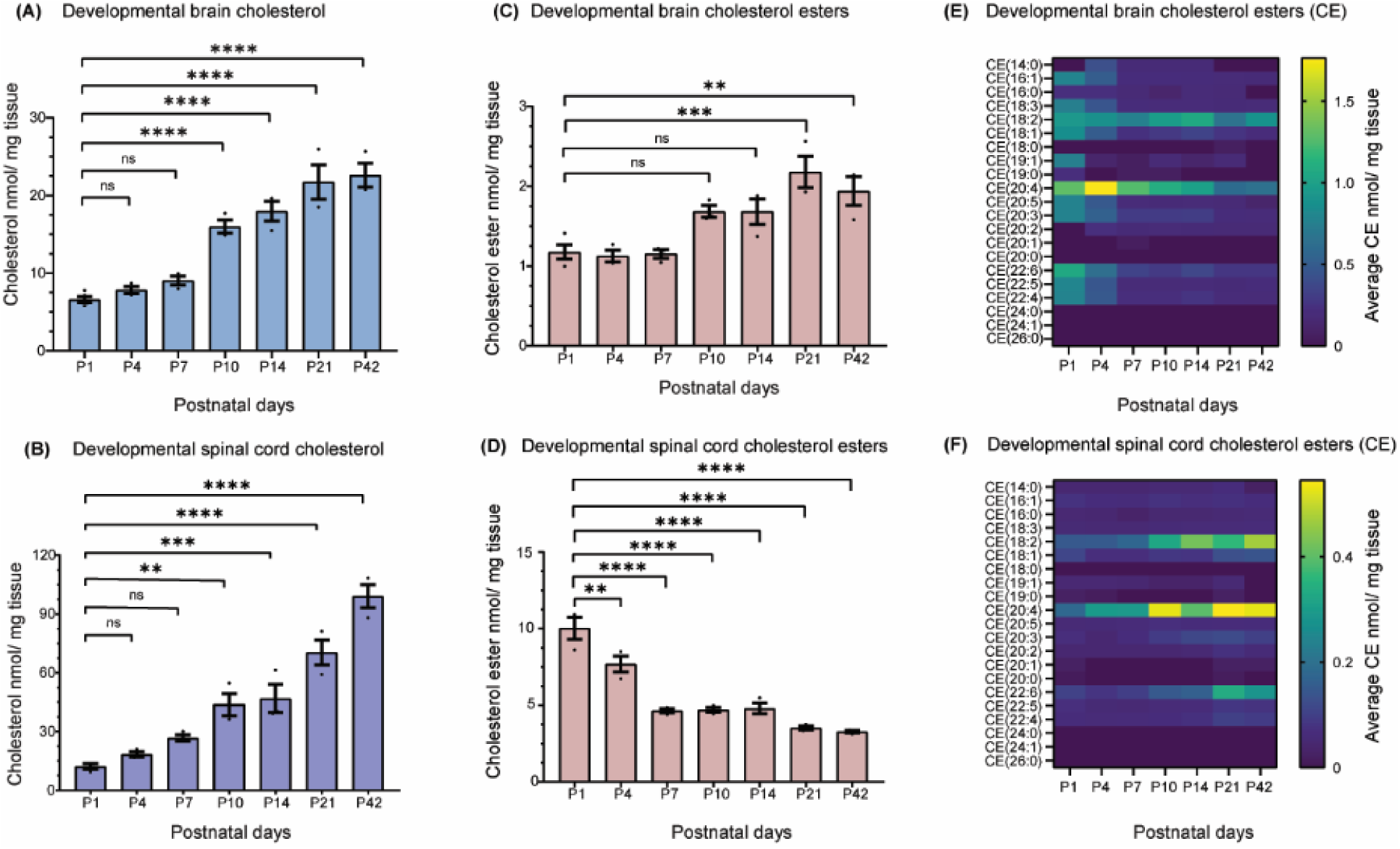
Cholesterol increases with age during development in the brain and spinal cord. **(A)**Total free cholesterol levels in the P1- P42 mouse brain. **(B)** Total free cholesterol levels in the mouse P1-P42 spinal cord. Ordinary one way-ANOVA, n= 3-5, Dunnett correction for multiple comparison data are plotted as mean ± SEM, GP: 0.123 (ns), 0.0332 (*), 0.0021 (**), 0.0002 (**), 0.0001 (***). **(C, D, E, F)** Cholesterol esters are accumulated during the early developmental spinal cord whereas it is accumulated at the later developmental time points in the brain. **(C)** Total cholesterol esters in the P1- P42 mouse brain. **(D)** Total cholesterol esters in the mouse P1-P42 spinal cord. Ordinary one way-ANOVA, n= 3-5, Tukey correction for multiple comparison data are plotted as mean ± SEM, GP: 0.123 (ns), 0.0332 (*), 0.0021 (**), 0.0002 (**), 0.0001 (***). **(E)** Heat map of the individual cholesterol ester type in the developmental brain (P1-P42). **(F)** Heat map of the individual cholesterol esters in the developmental spinal cord (P1-P42). Average amounts of each cholesterol ester in nmol/ mg tissue are shown in the heat maps.

We detected 21 cholesterol esters, with fatty acyl chains ranging from 14:0 to 22:6 in the postnatal mouse brain and spinal cord tissues (Supp. Table S7 & S9). Total cholesterol ester levels in the brain ranged from 1 nmol/mg to 3 nmol/mg and were stable across the time points measured (Figures 2C & 2E, Supp. Table S7). The spinal cord tissue was an average of 10 nmol/ mg cholesterol esters at P1 and decreased to 3 nmol/mg by P42 (Figure 2D & 2F, Supp. Table S9). The spinal cord had higher levels of total cholesterol esters as compared to the brain at developmental time points (8.6-fold higher than in the brain at P1 dropping to 1.8-fold by P42). In summary, total free cholesterol levels increased with developmental age in both the brain and the spinal cord, however, only the spinal cord had a high accumulation of cholesterol esters at P1 that rapidly decreased by P42.

### Cholesterol increases with aging in the brain, but not the spinal cord

In this study, we utilized a genetic mouse model of demyelination (*Plp1-CreERT; Myrf(fl/fl)*) to measure the dynamic changes of cholesterol over time in the brain and spinal cord (Figure 1B). Demyelination was induced with tamoxifen injections at 8 weeks of age, which caused ablation of *Myrf* (myelin regulatory factor) from mature oligodendrocytes. MYRF is a transcription factor that regulates the transcription of genes important in the production and maintenance of myelin (30). Deletion of *Myrf* resulted in oligodendrocyte cell death and gradual demyelination, however, *Myrf* was unaffected in oligodendrocyte precursor cells (OPCs). Following demyelination, OPCs proliferated and differentiated into mature myelinating oligodendrocytes leading to remyelination (29, 37). Clinically, *Myrf* ablation led to motor impairment measured by clinical scoring and rotarod analysis that was detectable by 5 weeks post-tamoxifen and peaked at 10-14 weeks post-tamoxifen. Motor symptoms gradually improved from 14 to 24 weeks post-tamoxifen (37, 38).

For the demyelination experiment, we collected tissues from 70-80 mice. TMS-derivatized cholesterol is only stable for ∼48 hours, which limited the number of samples that could be analyzed in a single batch for GC-MS cholesterol analysis. To overcome this problem, the samples were randomly divided into multiple batches to perform the lipid extraction, sample derivatization, and GC-MS analysis. Since GC-MS runs were performed on different days, separate internal standard calibration curves were prepared to avoid instrumental variations (LOD and LOQ for cholesterol 1.10 μM and 3.69 μM, respectively). To determine variation across batches, two-three quality control samples were analyzed with each batch. Quality control samples were prepared by mixing representative brain or spinal cord samples from each group. In Figure 3A, brain quality control samples show an average of 29 nmol/mg tissue concentration with a 12.9% coefficient of variation (CV), whereas it is 59 nmol/mg and 9.8 % CV for the spinal cord quality controls (Figure 3B).

**Figure 3:**
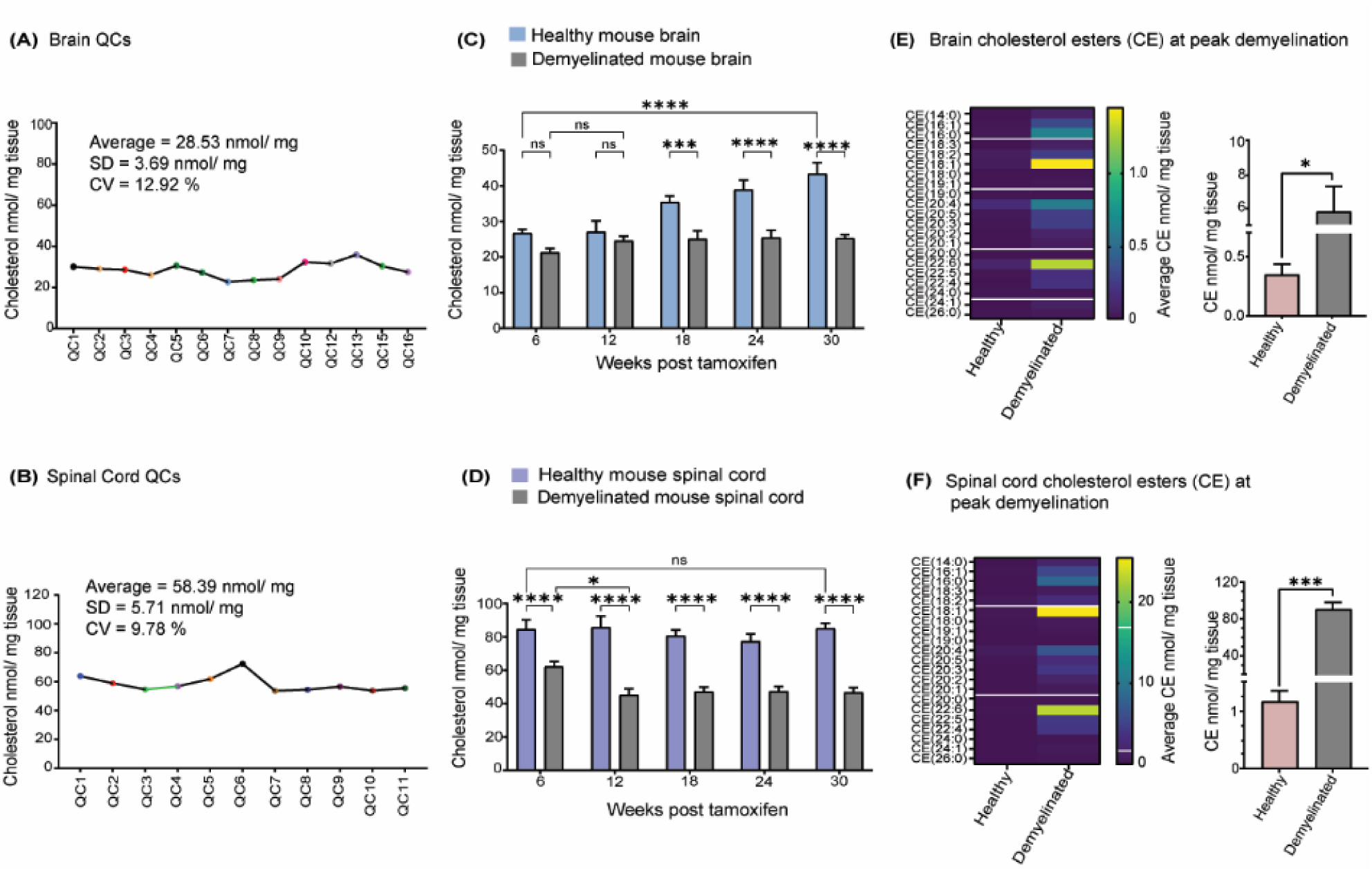
Total free cholesterol levels are reduced in both the brain and spinal cord in demyelinated mice. **(A)** Brain quality control samples (QCs) over the batches show an average of 28.53 nmol/ mg tissue with a coefficient variation of (CV) 12.92%, SD – Standard Deviation. **(B)** Spinal cord quality control samples (QCs) over the batches show an average of 58.39 nmol/ mg tissue with a coefficient variation of (CV) 9.78 %, SD – Standard Deviation. **(C)** Total free cholesterol levels in the *Plp* mouse brain. Two way-ANOVA, Holm-Šídák correction for multiple comparisons, n= 3-10, data are plotted as mean ± SEM, GP: 0.123 (ns), 0.0332 (*), 0.0021 (**), 0.0002 (**), 0.0001 (***). **(D)** Total free cholesterol levels in the *Plp* mouse spinal cord. Two-way ANOVA, Holm-Šídák correction for multiple comparisons, n= 3-10, data are plotted as mean ± SEM, GP: 0.123 (ns), 0.0332 (*), 0.0021 (**), 0.0002 (***), 0.0001 (****). **(E, F)** Cholesterol esters pool is increased at the peak demyelination time points of the brain and spinal cord. **(E)**Total amounts of cholesterol ester (CE) and individual CE types (heat map, CE nmol/ mg tissue) in the adult healthy and demyelinated mouse brain at week 12 post tamoxifen peak demyelination. **(F)**Total amounts of cholesterol ester (CE) and individual CE types (heat map, average CE nmol/ mg tissue) in the adult healthy and demyelinated mouse spinal cord at week 18 post tamoxifen peak demyelination. Unpaired two tailed t-test was performed for the bar graphs in **(E)** & **(F)**, CE data are plotted as mean ± SEM, GP: 0.123 (ns), 0.0332 (*), 0.0021 (**), 0.0002 (**), 0.0001 (***).

In healthy mouse brains, total free cholesterol was 27 nmol/mg tissue at 14 weeks of age (week 6 post-tamoxifen) and reached 43 nmol/mg tissue by 38 weeks of age (week 30 post-tamoxifen) (Figure 3C, Supp. Table S3). In the spinal cord, cholesterol levels remained relatively stable in the range of 77 nmol/mg to 86 nmol/mg tissue throughout all time points (14-38 weeks of age) (Figure 3D, Supp. Table S5). Thus, *de novo* cholesterol synthesis was still active in the brain in mice as old as 38 weeks, which is approaching “middle-aged” for a mouse (39, 40). In contrast, cholesterol levels in the spinal cord were at the same level from 14-38 weeks of age suggesting that cholesterol metabolism reached a steady state homeostasis by ∼3 months of age. At week 6 post tamoxifen, spinal cord cholesterol was three-fold higher than in the brain; however, as cholesterol increased with aging in the brain, the levels were only two-fold higher than the spinal cord at 30 weeks post-tamoxifen (38 weeks of age) (Figure 3C & 3D, Supp. Table S3 & S5). Even with the increase in brain cholesterol with aging, the spinal cord still had higher cholesterol consistent with higher myelin density as compared to the brain (2, 38).

### Brain and spinal cord free cholesterol levels are impaired with demyelination

The cholesterol levels in the brains of *Plp1*-iCKO-*Myrf* mice were measured during active demyelination (week 6 post tamoxifen), peak demyelination (week 12 post tamoxifen), initial recovery (week 18 post tamoxifen), and plateau recovery (weeks 24 and 30 post tamoxifen) (Figure 3C). Strikingly, the total free cholesterol did not drop significantly with the onset of demyelination at week 6 and remained consistent at ∼25 nmol/mg tissue during all subsequent time points, even during the remyelination period. Since the total free cholesterol levels in the healthy brains increased over that same time period, the brains at week 18, 24, and 30 post-tamoxifen had significant depletion in total cholesterol (Figure 3C, Supp. Table S3)(37). These results suggest that widespread demyelination interrupts the normal ongoing synthesis of CNS cholesterol that happens in healthy adult mice, and that even during robust remyelination at the later timepoints, the cholesterol synthesis rate did not recover to healthy levels.

In the spinal cord, cholesterol levels were significantly reduced in the demyelinated mice relative to healthy mice at all time points measured. Healthy spinal cord tissue at week 6 post-tamoxifen had an average of 85 nmol/mg tissue, and with demyelination, the cholesterol levels at week 6 dropped to 62 nmol/mg tissue and continued to decrease to the range of ∼45 nmol/mg tissue that persisted through the remaining time points (Figure 3D, Supp. Table S5). By the end of the experiment (week 30 post-tamoxifen), both the brain at 42% and the spinal cord at 45% showed similar levels of cholesterol depletion relative to healthy tissues (Supp. Table S3 & S5).

In our previous study, we measured the levels of cholesterol esters in the *Plp1*-iCKO-*Myrf* mice across the same time points examined here, and we observed major increases in cholesterol esters during peak demyelination over healthy tissues (30-fold in the brain and 130-fold in the spinal cord). In that study, cholesterol esters were measured by direct infusion of the sample into the mass spectrometer with no chromatographic separation. A semi-quantitative method was used based on internal standards spiked at known amounts, but a calibration curve was not prepared, and thus the amounts reported were not reflective of absolute levels. When we compared the cholesterol ester data obtained from our current quantitative method to the previous direct infusion method, we noticed that the levels of cholesterol esters detected in the healthy tissue in the developmental study were several fold higher than the healthy levels we previously reported suggesting that our previous measurements did not reflect the absolute levels. Thus, to obtain an absolute quantification of cholesterol esters during peak demyelination in the *Plp1*-iCKO-*Myrf* mice, we applied our new quantitative method to brain and spinal cord tissues from peak demyelination in the brain (week 12) and in the spinal cord (week 18). We observed healthy adult mouse brain had ∼0.3 nmol/ mg tissue total cholesterol esters, and the demyelinated mouse brain had elevated levels at ∼6 nmol/mg (16-fold increase). The healthy spinal cord had 1 nmol/mg cholesterol esters, which was dramatically elevated 77-fold to 91 nmol/mg during demyelination (Figure 3E and 3F, Supp. Table S11 & S12).

### Thyroid hormone action does not deplete total free cholesterol levels in the CNS

Sob-AM2 is a CNS-penetrating prodrug that can cross the blood-brain barrier to enter the central nervous system, where it is hydrolyzed to the parent thyroid hormone agonist known as sobetirome (41, 42). Sob-AM2 has previously been shown to improve remyelination and motor recovery in the *Plp1*-iCKO-*Myrf* demyelination model used in Figure 3 (37). In peripheral tissues, thyroid hormone is a well-established regulator of cholesterol metabolism and increased thyroid hormone lowers serum cholesterol through liver clearance mechanisms (43). Reduced cholesterol in the CNS is associated with impaired remyelination (7, 44), and therefore, we tested how Sob-AM2 affected total cholesterol levels in the brain and spinal cord.

Mice were treated with chow containing Sob-AM2 starting 2 weeks post-tamoxifen. Tissue samples were analyzed from week 12 post tamoxifen (peak demyelination) and week 18 post tamoxifen (recovery). Overall, the mice treated with SobAM2 showed similar overall trends to the vehicle mice with reduced cholesterol levels in demyelinated mice as compared to healthy mice in both brains (Figure 4A, Supp. Table S4) and spinal cords (Figure 4B, Supp. Table S6). No differences were observed due to drug treatment. Considering the critical role of cholesterol in CNS myelination and remyelination, our observation that thyroid hormone agonists do not reduce brain cholesterol in healthy brains or chronically in the context of demyelination is a positive indication for the therapeutic potential of Sob-AM2 as a remyelinating agent.

**Figure 4:**
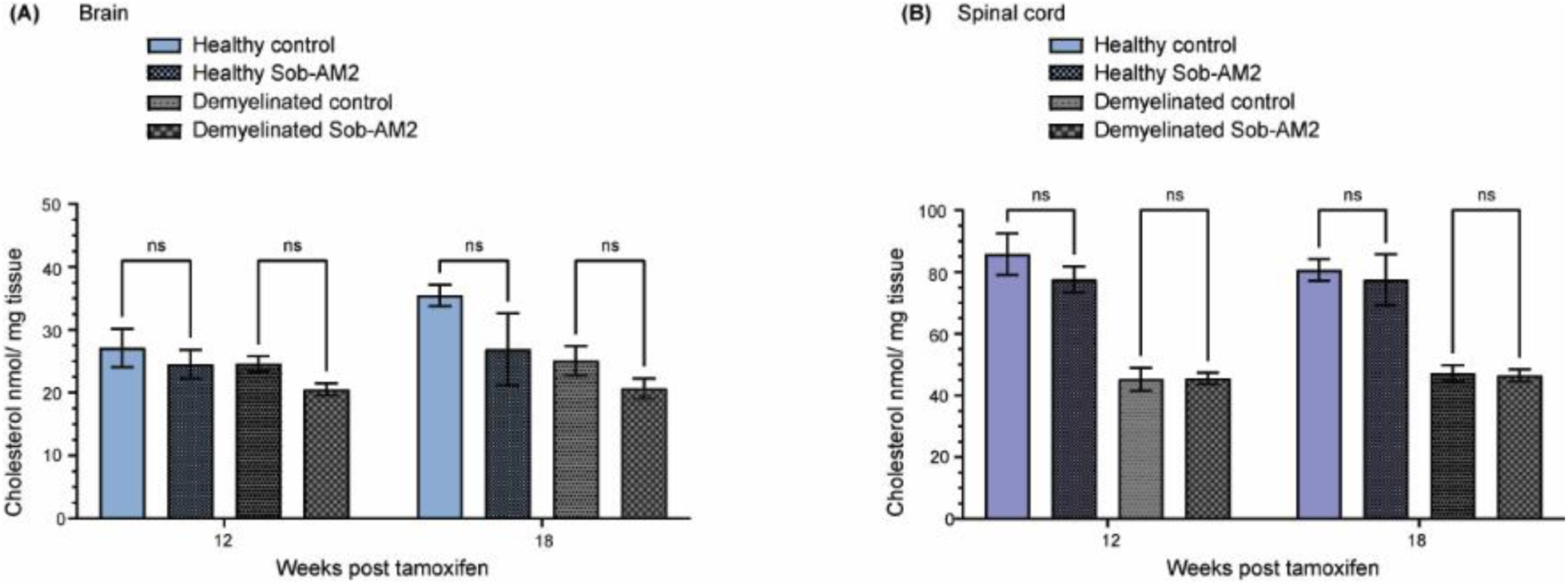
Sob-AM2 enhances remyelination without significantly affecting CNS total cholesterol levels. **(A)** Effect of Sob-AM2 drug on cholesterol levels in the brain. **(B)** Effect of the Sob-AM2 drug on cholesterol levels in the spinal cord. Two-way ANOVA (mixed models, Tukey correction for multiple comparisons n=4-10, data are plotted as mean ± SEM, GP: 0.123 (ns), 0.0332 (*), 0.0021 (**), 0.0002 (**), 0.0001 (***).

## 5. DISCUSSION

### *de novo* cholesterol synthesis in the brain

The brain is considered the most cholesterol rich organ (15 – 20 mg/g tissue) containing 20% of the whole-body cholesterol (3, 45). Most of brain cholesterol (about 70% - 80%) is in the myelin sheath, where it is an essential structural component, but cholesterol also serves as a precursor of steroid hormones (46) and a required component of synapses (11). Brain cholesterol metabolism is separated from peripheral cholesterol metabolism by the blood-brain barrier (45). Brain cholesterol is primarily generated by astrocytes and is known to be transported by apolipoprotein E (apoE) to neurons (47). However, both oligodendrocytes and neurons are also capable of *de novo* synthesis of cholesterol (7, 22). Brain cholesterol is known to accumulate during the prenatal period and adolescence correlating with the increase in myelin levels. In the adult brain, cholesterol has low turnover with a long half-life (3). Excess cholesterol in the brain is esterified (∼1%) and stored as lipid droplets inside cells (8), or converted to oxysterols that can cross the blood-brain barrier (48).

### Cholesterol synthesis during brain and spinal cord development and maturation

Myelination begins in the rodent brain around P10, and mass of the brain doubles and water content decreases during this phase. Following the initial rapid phase of myelination, slower myelination continues during the final brain maturation stages. Previous studies have suggested that cholesterol levels track with myelination. Cholesterol in the developing rat brain was 10.7 μmol/ g wet weight (P3), 12.6 μmol/ g (P6), and 22.6 μmol/ g (P12) (49). It has also been observed that rates of cholesterol synthesis in the CNS were high (app. 0.26 mg/day) in the CNS during first three weeks after birth, and the rate declined to a constant value (app. 0.035 mg/ day) in older mice at 13 -26 weeks old (50). The total cholesterol levels in our data align with the brain developmental phases and the cholesterol levels previously described. We observed that cholesterol levels increased rapidly at P10 and then gradually increased until P42 in both the brain and spinal cord (Figure 2 A & B, Supp. Tables S1 & S2). In humans, prolonged myelination starts prenatally in humans whereas it mainly occurs postnatally in rodents (51). Plasma cholesterol levels correlate between the human fetus and the mother at the fetus ages younger than 6 months, but after that point the fetus produces the majority of cholesterol by itself (52). However, the specific time where the blood brain barrier becomes not permeable to cholesterol during brain development in humans has not been determined yet (51).

Following the developmental phase, we observed that total brain cholesterol levels in healthy mice increased from 14 to 38 weeks of age (Figure 3C, Supp. Table S3), which is consistent with previous measurements of brain cholesterol (50, 53). Increasing cholesterol during adulthood also correlates with the ongoing myelination in the corpus callosum and cortex of mice that continues through adulthood (54–56). In contrast to the brain, our spinal cord data showed that cholesterol increased during development (up to P42) and then stabilized 6-30 weeks post-tamoxifen (14-38 weeks of age) (Figure 2B and 3D, Supp. Tables S2 & S5). During nervous system development and myelination, the spinal cord is fully myelinated before the brain, and thus spinal cord cholesterol dynamics that we measured correlate with known myelination trends (57).

During development, the brain initially (P1-P7) had ∼1 nmol/mg tissue of cholesterol esters (∼15% of total pool), which increased to 1.6-2.0 nmol/mg levels by P10 (∼7% of total pool). Since the free cholesterol levels continuously increased in the postnatal mouse brain, the percentage of cholesterol esters in the total cholesterol pool decreased with age (Figure 5A, Supp. Table S8). Notably, the spinal cord has a very high accumulation of cholesterol esters (10 nmol/ mg at P1, ∼45% of total pool) at early postnatal time points which decreased to 3.0 nmol/mg by P42 (∼3% of total pool) (Figure 5B, Supp. Table S10). The high level of cholesterol esters during early development could indicate that there is a large pool of excess cholesterol in spinal cord, which may be related to the high demand for cholesterol at this time point. In healthy brain and spinal cord during adulthood, cholesterol esters make up 1.0 −1.5% of the total cholesterol pool.

**Figure 5:**
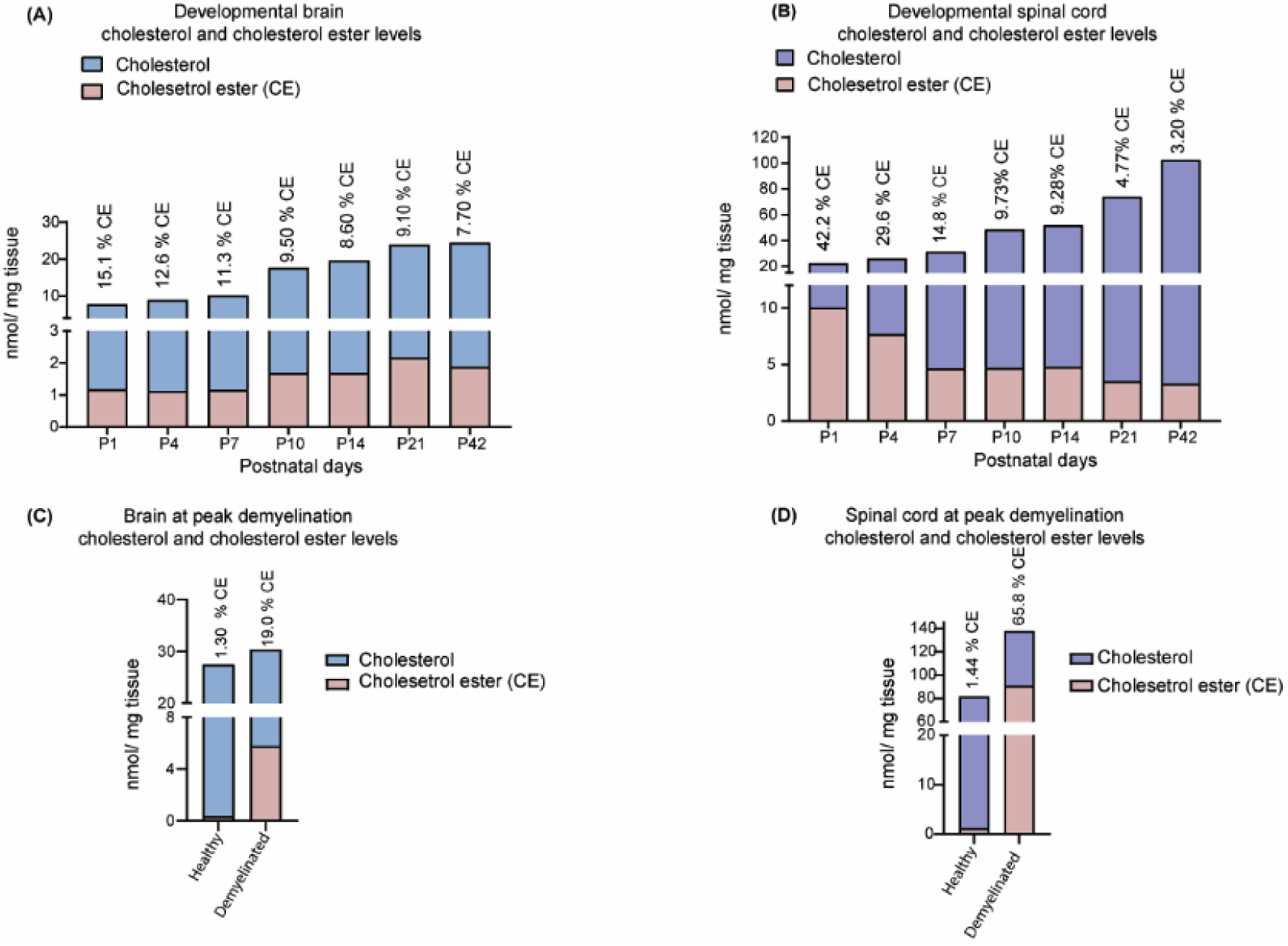
Percentage of cholesterol esters in the total cholesterol pool is decreasing during the development in both brain and spinal cord. **(A)** Calculated CE percentage the total cholesterol pool in the developmental brain during P1 -P42. **(B)** Calculated CE percentage and the total cholesterol pool in the developmental spinal cord during P1 -P42. Brain and spinal cord tissue show higher percentage of CE accumulation at the peak demyelination compared to healthy CNS tissues. **(C)** Healthy and demyelinated brain tissue total cholesterol pool and the percentages CE. **(D)** Healthy and demyelinated spinal cord tissue total cholesterol pool and the percentages CE. Average cholesterol and cholesterol levels are plotted in the combined bar graphs above. Please refer to Figures 2 and 3 for all individual data included with error bars and statistics.

### Brain and spinal cord cholesterol levels drop during demyelination

The pathology of multiple sclerosis is also associated with altered cholesterol and cholesterol metabolite biomarkers in the serum or cerebrospinal fluid (21, 58), however, it still remains unclear whether peripheral dyslipidemia (high cholesterol and lipids) causes multiple sclerosis pathology or vice versa (59). In human MS lesion samples, cholesterol synthesis gene expression was decreased in astrocytes and neurons, but increased in oligodendrocytes (22), suggesting cell-specific responses to inflammatory demyelination. In the brains of *Plp1*-iCKO-*Myrf* mice undergoing demyelination, we observed only a small loss of cholesterol at the initial timepoints (week 6 and week 12 post tamoxifen). However, during the remyelination phase, total cholesterol levels remained steady (∼25 nmol/mg) and did not increase with age as occurred in healthy mice brains (Figure 3C, Supp. Table S3). This result was somewhat surprising as both histological and motor testing have demonstrated increased myelin levels and improved motor performance at weeks 18 and 24 (38), which might be expected to correlate with increased cholesterol production and may suggest that the observed remyelination occurs primarily through cholesterol recycling. Further, the data suggests that the widespread demyelination in the *Plp1*-iCKO-*Myrf* model disrupts ongoing normal cholesterol synthesis that occurs during healthy aging.

Our results also contrast with previous findings using chronic cuprizone administration in mice that showed increased levels of cholesterol, desmosterol, and lathosterol in isolated neurons during remyelination as compared to the healthy controls (22), however, total levels across the brain were not measured and we did not examine cell-specific levels. The cuprizone model also features more localized demyelination than the *Plp1*-iCKO-*Myrf* model in which all oligodendrocytes are affected, which may lead to model-specific differences in both cellular and global cholesterol homeostasis. Cholesterol synthesis is controlled by a complicated feedback mechanism involving transcriptional, translational, and post-translational regulation (60) and future research will focus on identifying which mechanisms are involved in regulating cholesterol biosynthesis in the brain after demyelination.

In contrast to the brain, the spinal cord had a much larger drop in total cholesterol during the initial demyelination phase (∼26 % loss). Following the initial drop, the spinal cord reached a steady plateau level during all time points (∼45 nmol/ mg tissue) (Supp. Tables S3 & S5). Since the spinal cord cholesterol levels do not appreciably increase after 14 weeks of age, it is not surprising that total cholesterol did not increase after the primary demyelination insult. The drop in cholesterol with no subsequent increase is consistent with the limited remyelination observed in the spinal cord of this model (38).

### Trends of total free cholesterol and cholesterol ester in demyelinated mice

Although cholesterol esters are not highly abundant in healthy CNS tissue (<1% of total cholesterol) (8), accumulation of cholesterol esters has been observed in postmortem CNS tissues from patients with myelin damage (23), and in cerebrospinal fluid of MS patients with more severe symptoms (24). Cholesterol esters also accumulate in the CNS in animal models of demyelination (23). Recently, it was established that cholesterol ester levels increased during demyelination and decreased with remyelination in the *Plp1*-iCKO-*Myrf* model (28). To analyze how the overall cholesterol pool is changing in the model, we compared the cholesterol and cholesterol ester levels in the brain and spinal cord during peak demyelination .At peak demyelination we observed that 1.3% of CE percentage in the total cholesterol pool observed in healthy mice goes up to 19% higher with demyelination in the brain (Figure 5C, Supp. Table S11). Interestingly, the adult healthy mouse spinal cord also has a similar range of total cholesterol ester levels (1 nmol/mg) accounting 1.4% of the total cholesterol pool. However, with demyelination, the total cholesterol esters go higher as 66% at the peak demyelination (Figure 5D, Supp. Table S12). This indicates a substantial flux of cholesterol through the cholesterol ester pathway and a large potential pool of cholesterol that could be stored for later remyelination. However, cholesterol ester formation explains part of the loss of the cholesterol during demyelination, but it does not explain the chronic reduction in brain cholesterol biosynthesis.

The chronic cholesterol loss that we observe during demyelination in the *Plp1*-iCKO-*Myrf* model likely occurs through the conversion of cholesterol into oxysterol and other oxidative byproducts. Cholesterol in myelin debris is phagocytosed by glial cells, leading to the local downregulation of cholesterol synthesis and increased synthesis of some oxysterols (44). In particular, 24S-hydroxycholesterol can pass through the blood-brain barrier and will not be detected in the brain homogenate (48). Detailed mass spectrometry analysis of cholesterol synthesis pathway intermediates, cholesterol metabolites, and oxysterols will be required to discern how cholesterol biosynthesis is affected in the model and whether cholesterol is converted to other metabolites during demyelination.

In this study, we measured how brain and spinal cord cholesterol levels fluctuate during postnatal myelination, demyelination, and remyelination in an inducible genetic mouse model using an internal standard-based quantification of cholesterol with GC-MS-SIM and cholesterol esters with LC-MS. Cholesterol in the healthy mouse brain increased steadily up to 38 weeks while cholesterol in the spinal cord increased during developmental phases (up to P42) and then remained steady through 38 weeks of age. Cholesterol esters were relatively high at P1 in the spinal cord (∼40% of total pool), but levels dropped to ∼1% in both the brain and spinal cord during development and adulthood. Both the demyelinated mouse brain and the spinal cord had significant reductions in unesterified cholesterol during the remyelination phase (28), and cholesterol never returned to healthy levels. The longitudinal cholesterol data indicates that normal cholesterol biosynthesis is disrupted in the brain after demyelination in this model and that cholesterol depletion persists in the spinal cord. Moreover, there is a large accumulation of cholesterol esters in the total cholesterol pool (∼20% in the brain and ∼60 % in the spinal cord) during peak demyelination whereas it is only ∼1% in both healthy brain and spinal cord. Future studies will focus on revealing the mechanisms behind the chronic loss in cholesterol levels. Identifying targets to preserve CNS cholesterol or restore cholesterol biosynthesis could be a promising therapeutic strategy for neurodegenerative diseases and age-related myelin degeneration.

## Supporting information

Manuscript Supplementary Table S1-S12

## 6. DATA AVAILABILITY STATEMENT

All analyzed mass spectrometry data is available with the manuscript or the supplementary information. Raw mass spectrometry files will be available in the Metabolomics Workbench repository.

## 7. ACKNOWLEDGEMENTS

Gas chromatography-mass spectrometry was performed in the Theodore “Ted” Kuwana Shimadzu Precision Instrumental Laboratory in the Department of Chemistry at the University of Kansas.

## 9. AUTHORS CONTRIBUTIONS

Disni Dedunupitiya: Methodology, Formal analysis, Investigation, Writing – Original Draft, Writing – Review & Editing, Visualization.

Eden P. Go: Methodology, Investigation, Formal Analysis, Writing – Review & Editing.

Travis Witte: Methodology, Resources.

Ashlynn Elliott: Investigation.

Nishama De Silva Mohotti: Investigation.

Jenna M. Williams: Investigation.

Hiroko Kobayashi: Investigation.

Rashmi Binjawadagi: Investigation.

Heather Desaire: Methodology, Resources, Writing – Review & Editing.

Meredith Hartley: Conceptualization, Methodology, Writing – Review & Editing, Supervision, Project administration, Funding acquisition.

## 10. FOOTNOTES TO TEXT (IF ANY)

## 11. SUPPLEMENTARY TABLES

Supplementary Table S1: Developmental free cholesterol level in the mouse brain P1- P42.

Supplementary Table S2: Developmental free cholesterol level in the mouse spinal cord P1- P42.

Supplementary Table S3: Free cholesterol level in the adult healthy and demyelinated mouse brains.

Supplementary Table S4: Free cholesterol level in the adult mouse brains with Sob-AM2 drug treatment.

Supplementary Table S5: Free cholesterol level in the adult healthy and demyelinated mouse spinal cords.

Supplementary Table S6: Free cholesterol level in the adult mouse spinal cords with Sob-AM2 drug treatment.

Supplementary Table S7: Cholesterol ester amounts in the developmental brain P1-P42.

Supplementary Table S8: Cholesterol ester percentage over total cholesterol pool in the developmental brain P1-P42.

Supplementary Table S9: Cholesterol ester amounts in the developmental spinal cord P1-P42.

Supplementary Table S10: Cholesterol ester percentage over total cholesterol pool in the developmental spinal cord P1-P42.

Supplementary Table S11: Cholesterol ester amounts in healthy and demyelinated brain at peak demyelination.

Supplementary Table S12: Cholesterol ester amounts in healthy and demyelinated spinal cord at peak demyelination.

